# Engineering programmable CAR-T cells with tunable controls as fail-safe mechanisms during cancer immunotherapy

**DOI:** 10.1101/2024.07.09.602762

**Authors:** Mikail Dogan, Lina Kozhaya, Lindsey Placek, Ece Karhan, Mesut Yigit, Christina Mohammed, Qi Zhao, Derya Unutmaz

**Author notes:** These authors contributed equally (Mikail Dogan, Lina Kozhaya).

## Abstract

Chimeric antigen receptor (CAR) T-cell therapies have demonstrated remarkable efficacy in treating various cancers, but significant risks of severe adverse effects remain. In this study, we present a synthetic biology toolbox for engineering CAR-T cells with tunable and controllable features to mitigate these risks. Using this toolbox, we programmed cytotoxic CAR-T cells to be turned on and off using synthetic genes with tetracycline response elements, referred to as iOnCAR and iOffCAR. We achieved temporal regulation of CAR expression using the synthetic Notch receptor system and “AND-gate” programs, which allow for additional control using doxycycline. We also engineered cooperation among T cells through CD4 help to CD8 T cells in targeting cancer cells via synthetic Notch receptors controlled by doxycycline treatment. Additionally, we developed a T cell fratricide system by triggering a genetic “Battle Royale” mechanism that enables CAR-T cells to eliminate each other. In this system, surviving T cells become activated, proliferate, and display high cytotoxicity to target cells. In conclusion, our synthetic biology toolbox provides targeted fail-safe solutions for improving the safety and potency of CAR-T cell therapies by integrating different genetic circuits with a commonly used antibiotic.

## Introduction

The development of chimeric antigen receptor (CAR) T-cell therapies has revolutionized the treatment of various cancers, especially hematological malignancies that are refractory to conventional therapies. Many clinical trials have demonstrated the remarkable efficacy and durability of CAR-T cell therapies in patients with B cell malignancies, and some studies report promising results in solid tumors^1–4^. However, these therapies also pose significant risks of severe adverse events, such as cytokine release syndrome (CRS), neurotoxicity, and on-target off-tumor toxicity^5–8^.

Several strategies have been developed to modulate CAR expression or activity to enhance CAR-T cell therapies’ safety and versatility^9–11^. One such strategy introduced “suicide switches” that induce apoptosis in CAR-T cells in response to exogenous molecules^12–16^. This allowed for the rapid elimination of the engineered T cells in case of severe toxicity. However, suicide genes are designed to eliminate the engineered T cells irreversibly, which compromises long-term protection against the original target and undermines the considerable time, effort, and expense involved in developing the therapy. Furthermore, the exogenous molecules used in these studies to activate suicide genes may have undesirable side effects or low bioavailability. For instance, some suicide switches require immune-suppressant small molecules that may increase the risk of infection or malignancy^15,17^.

Other studies have used inducible promoters to control CAR expression by exogenous molecules that act as transcriptional activators, such as the Tet-on system^18–20^. This allows for flexible adjustment of CAR-T cell function depending on the clinical situation. However, this system requires continuous administration of the stimulant that may have undesirable side effects. Synthetic Notch receptors (synNotch) have also been used for fail-safe purposes to induce CAR expression or activity by a secondary antigen on target cells^21–23^. These are artificially designed receptors that consist of an extracellular domain that recognizes a specific ligand and an intracellular domain that activates a transcriptional program upon ligand binding. By coupling synNotch receptors with CARs, researchers achieved antigen-specific activation of CAR-T cells only when the CAR-T cells encounter tumor cells that express both antigens^21–23^. Nevertheless, synNotch receptors may benefit from additional regulation mechanisms to fine-tune antigen-specific activation.

In this study, using genetic programming, we developed a set of CAR-T control tools. We developed a tetracycline response element inducible system using doxycycline (dox) for the induction or suppression of CAR expression, allowing flexible adjustment of CAR-T cell function in response to tumor cell activation. This system also modulated synNotch receptor-mediated CAR induction triggered by target antigens, enabling fine-tuning of specific activation and reducing the risk of on-target off-tumor toxicity. Additionally, we developed a T cell cooperative approach by engineering CD4+ T cells to express a synNotch receptor ligand that induces CAR expression in CD8+ T cells. Furthermore, we created a novel self-kill switch we called “battle royale”, which induces fratricide within CAR-T cells, both expanding and improving the cytotoxic efficiency of the surviving T cells. Together, these synthetic biology approaches have the potential to be important fail-safe tools in engineering CAR-T cells with tunable and controllable genetic circuits for clinical applications in cancer immunotherapy.

## Results

To rapidly and reliably test the fail-safe approaches for CAR-T cells, we utilized a flow cytometry-based cytotoxicity assay that measures the killing of CD19+ target cells by anti-CD19 CAR-engineered primary human T cells (Fig. 1a). Briefly, a CAR cassette, described in the methods was cloned into lentiviruses (Fig. 1b). These lentiviruses were then used to transduce primary CD8+ T cells to constitutively express these CARs on their surface. An empty vector lentivirus was transduced into CD8+ T cells as a control. After 10-12 days of cell expansion, we assessed the CAR expression on the primary T cells using a recombinant CD19-Fc protein and an anti-Fc antibody. CD8+ T cells engineered with CAR-expressing lentiviruses displayed strong surface expression of the construct, while the empty vector control only showed the GFP marker with no surface CAR detection (Fig. 1c).

**Figure 1.**
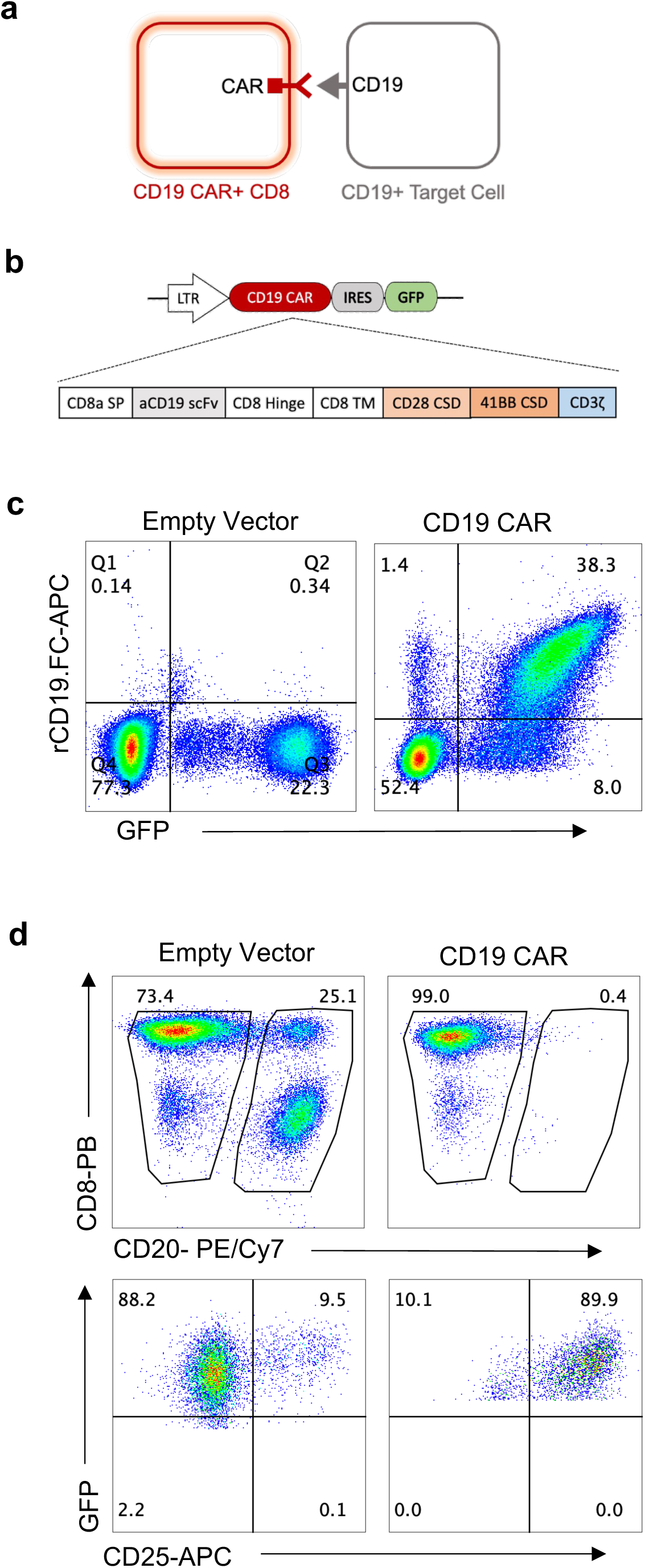
Generation and characterization of CD19-specific CAR-T cells. **a,** Illustration of anti-CD19 CAR-expressing CD8+ T cells interacting with CD19-expressing target cells. **b,** Schematic representation of anti-CD19 CAR construct co-expressed with a GFP reporter via an Internal Ribosomal Entry Site (IRES) under LTR promoter. The CAR construct comprises of CD8 alpha signal peptide, single chain variable fragment (scFv) of a CD19, CD8-derived hinge and transmembrane domains, 4-1BB (CD137) co-stimulatory domain, and a CD3ζ domain. **c,** Flow cytometry plots demonstrating the expression of CD19 CAR construct on CD8+ T cells. Isolated CD8+ T cells were activated and transduced with lentiviruses encoding CD19 CAR and empty vector constructs. After 10-12 days of culture, the cells were stained with CD19-Fc fusion protein and anti-human Fc secondary antibody to assess surface expression of anti-CD19 CAR, which was correlated with the GFP reporter. **d**, Representative experiment demonstrating the cytotoxicity of CD19 CAR CD8+ T cells targeting CD19+ T2 cells. CD8+ cells transduced with an empty vector were used as a control. Effector and target cells were identified with CD8 and CD20 surface expression, respectively. CD25 expression was used as a marker to evaluate T-cell activation. The experiments were replicated twice at different time points (n=3).

Next, we performed a cytotoxicity assay by co-culturing CAR-T cells with a CD19+ B cell line, T2 cells. On day 2, the cells were stained with CD8 and CD20 antibodies to identify effector and target cells, respectively, and with a CD25 antibody to evaluate the activation of the effector cells (Fig. 1d). CAR-T cells effectively eliminated their CD19+ targets and exhibited high levels of T cell activation based on upregulation of CD25 expression (Fig. 1d).

We next constructed a simple on/off switch at the transcription level of CAR expression in primary T cells using tetracycline-on and tetracycline-off (tet-on and tet-off) systems, as this is a well-established method for conditional gene expression regulation in mammalian cells^24^. Our goal was to engineer fail-safe CAR-T cells capable of regulated expression of inducible CARs (iOnCARs), such that cells would only exhibit cytotoxicity in the presence of doxycycline, a more stable tetracycline analog^25^, and the target antigen, thus creating an AND-gate operation at the cellular level (Fig. 2a).

**Figure 2.**
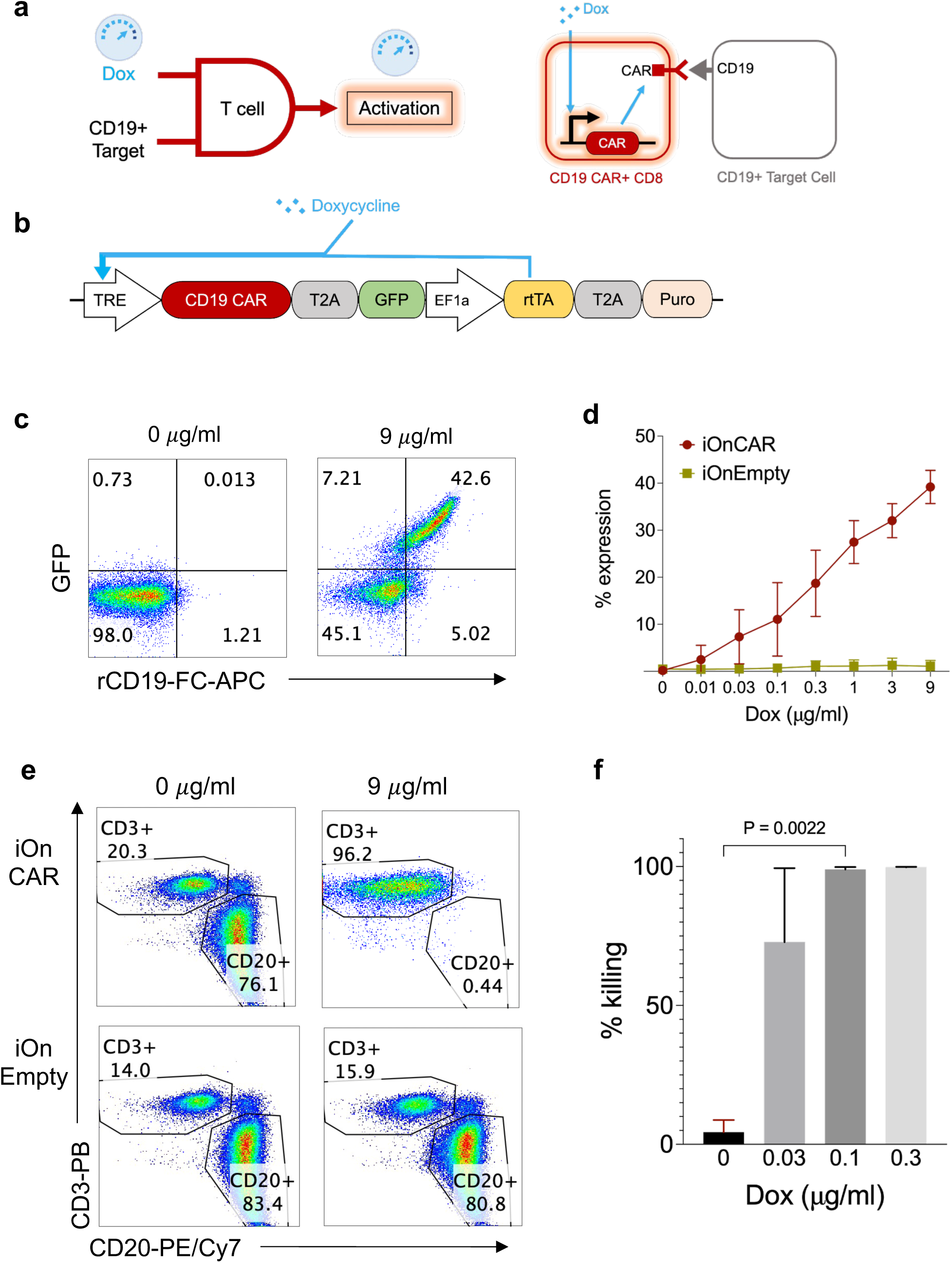
Doxycycline-inducible CAR-T cells: design, expression and cytotoxicity. **a,** Illustration of engineered T cells becoming activated upon interacting with CD19+ target cell in the presence of doxycycline (dox) in a tunable manner (left) and doxycycline-inducible CAR-T (iOnCAR) cell interacting with CD19+ target cell (right). **b**, Schematic outline of the iOnCAR lentiviral vector construct. EF1 promoter driving reverse tetracycline-controlled transactivator fusion protein (rtTA) to control Tet-inducible TRE promoter driving CAR-IRES-GFP in the presence of dox. The puromycin resistance gene (Puro) is also expressed under the EF1 promoter via the T2A cleavage site (T2A). **c**, Representative flow cytometry plots showing the iOnCAR expression on primary CD8+ T cells in the absence and presence (9 *μ*g/ml) of dox. **d**, Dose-dependent CAR expression 48 hours after the addition of dox. CD8+ T cells transduced with empty vectors were used as controls. The error bars represent mean and one standard deviation. **e**, Cytotoxicity of iOnCAR CD8+ T cells targeting CD19+ T2 cells in the absence and presence (9 *μ*g/ml) of dox. CD8+ T cells transduced with empty vectors were used as controls. Effector and target cells were identified with CD8 and CD20 surface expressions, respectively. **f**, Cytotoxicity of iOnCARs targeting CD19+ T2 cells in the absence and presence of dox titrated from 0.03 *μ*g/mL to 0.3 *μ*g/mL in a 3-fold manner. The percent cytotoxicity was calculated as described in the methods using the percentage of CD20+ cells in the condition where iOnEmpty T cells were combined with T2 cells. The error bars represent mean and one standard error of mean. The experiments were replicated twice at different time points (n=3). A paired t-test was used to determine the statistical significance.

The iOnCAR construct was designed to be integrated into the original CAR construct (Fig. 2b). To test the expression of iOnCAR, CD8+ T cells transduced with lentiviruses carrying the construct were assessed after 10-12 days of in vitro expansion. An empty vector (iOnEmpty) was used to transduce primary CD8+ T cells to generate control cells. The CAR-T cells were then treated with doxycycline (dox) at different concentrations ranging from 0.01 ug/mL to 9 ug/mL and were stained for CAR expression. Treatment with 9 ug/mL induced robust CAR expression in cells co-transduced with iOnCAR and a GFP marker, whereas control cells were double negative (Fig. 2c). The strength of CAR expression correlated with dox concentration, suggesting fine-tuned control of expression (Fig. 2d). Next, iOnCARs and CD19+ target cells were combined in a 1:3 E:T ratio in the presence and absence of dox for 3 days to determine cytotoxic capacity. Cytotoxicity assays revealed that iOnCAR CD8+ T cells efficiently eliminated their CD19+ target cells compared to control cells (Fig. 2e). After three days of incubation, cytotoxicity plateaued at nearly 100% killing in the presence of 0.1 ug/mL dox (Fig. 2f).

One caveat of the inducible tet-on approach is that it requires the continuous presence of dox. Therefore, we also explored the reverse approach and integrated the tet-off system into CAR-T cells. The cytotoxicity of “iOffCAR” T cells would require the absence of dox and the presence of target cells; hence, their native state would be cytotoxic, but cytotoxic activity could be turned off with dox treatment, if clinically warranted (Fig. 3a). We swapped the tet-on transcription factor (rtTA) with the tet-off transcription factor (tTA) to render the construct repressible by dox (Fig. 3b). To test the effectiveness of the system, we treated the transduced CAR-T cells with 1 ug/mL dox for six days and stained them for CAR expression. The dox treatment of iOffCAR cells greatly reduced CAR expression (Fig. 3c). To assess the rate of CAR downregulation, we performed a time course experiment and determined that dox treatment reduced CAR expression by roughly 50% every 2 days and reached less than 1% of initial expression after 6 days (Fig. 3d). Subsequent cytotoxicity assays showed that iOffCAR cells treated with 1 ug/mL dox were highly effective at eliminating target cells, while iOffCAR cells without dox displayed no cytotoxicity, similar to the CD8+ T cells engineered with iOffEmpty lentiviruses (Fig. 3e, f).

**Figure 3.**
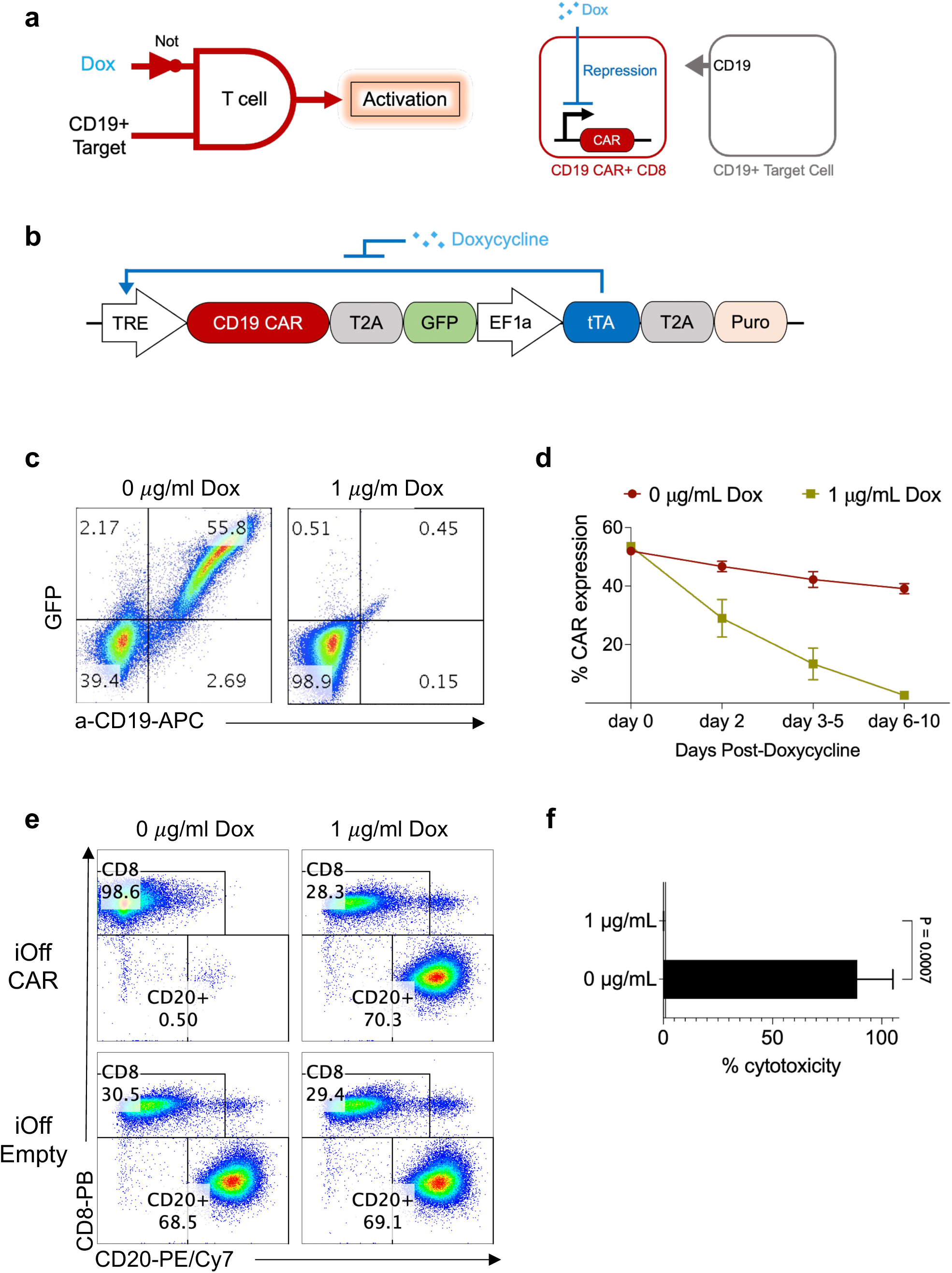
Developing doxycycline-repressible CAR-T cells. **a**, Illustration of engineered T cells activating upon interacting with CD19+ target cell in the absence of doxycycline (dox) (left) and dox-repressible CAR-T (iOffCAR) interacting with CD19+ target cell (right). **b**, Schematic outline of the iOffCAR lentiviral vector construct. EF1 promoter driving tetracycline transactivator fusion protein (tTA) to switch off Tet-inducible TRE promoter driving CAR-IRES-GFP when dox is present. The puromycin resistance gene (Puro) is also expressed under the EF1 promoter via the T2A cleavage site (T2A). **c**, iOffCAR T cells losing their CAR expression in the presence of dox (1 *μ*g/ml). **d**, CAR expression of iOffCAR T cells in the absence and presence of dox in a time-dependent manner. The error bars represent mean and one standard error of mean. **e**, Cytotoxicity of iOnCAR CD8 T cells targeting CD19+ T2 cells in the absence and presence (1 *μ*g/ml) of dox. CD8+ T cells transduced with empty vectors were used as controls. Effector and target cells were identified with CD8 and CD20 surface expressions, respectively. **f**, Cytotoxicity of iOffCARs targeting CD19+ T2 cells in the absence and presence of dox (1 *μ*g/mL). The percent cytotoxicity was calculated as described in the methods using the percentage of CD20+ cells in the condition where iOffEmpty T cells were combined with T2 cells. The error bars represent mean and one standard deviation. The experiments were replicated twice at different time points (n=3). An unpaired t-test was used to determine the statistical significance.

The synthetic notch (synNotch) system uses a synthetic receptor to sense a specific ligand on the target cell surface and control downstream gene expression^23,26,27^. We sought to incorporate this into our CAR-T cell fail-safe toolbox, envisioning a system where a marker on the target cell would induce CAR expression, and in turn, trigger a second marker on the target cell to induce apoptosis, adding additional control to our AND-gate fail-safe mechanism (Fig. 4a). We engineered an anti-GFP synNotch receptor into our CAR-T cells that would sense surface GFP as a ligand and release a tTA transcription factor, which would induce a TRE promoter to turn on the TRE-CAR construct. Since tTA is sensitive to dox, this system can be regulated with this molecule as an extra layer of control by modulating the binding affinity of tTA to the TRE promoter (Fig. 4a). In the presence of dox, tTA could not bind to the TRE promoter and activate the expression of the CAR construct, thus preventing the cytotoxicity of synNotch CD8+ T cells against GFP+ T2 cells. Accordingly, we transduced primary CD8+ T cells with lentiviruses encoding the anti-GFP-tTA synNotch receptor and the TRE-CAR construct constitutively expressing mCherry as a reporter and expanded them for 14 days. Additionally, we engineered GFP-expressing T2 cells that naturally express CD19 as target cells. Wild-type CD8+ T cells and wild-type T2 cells were used as controls. We performed a cytotoxicity assay by (I) co-culturing wild-type CD8+ T cells with GFP+ T2 cells, (II) synNotch CD8+ T cells with wild-type T2 cells, (III) synNotch CD8+ T cells with GFP+ T2 cells in the absence or (IV) presence of dox for 3 days (Fig. 4b). The cytotoxicity assay revealed that only synNotch CD8+ T cells, in the absence of dox, upgregulated CAR expression and could eliminate their GFP+ T2 target cells (Fig. 4c, d, panel III). Addition of dox switched off CAR expression induced by Notch and thus functioned as a further control layer on the synNotch receptor system (Fig. 4b, c, panel IV).

**Figure 4.**
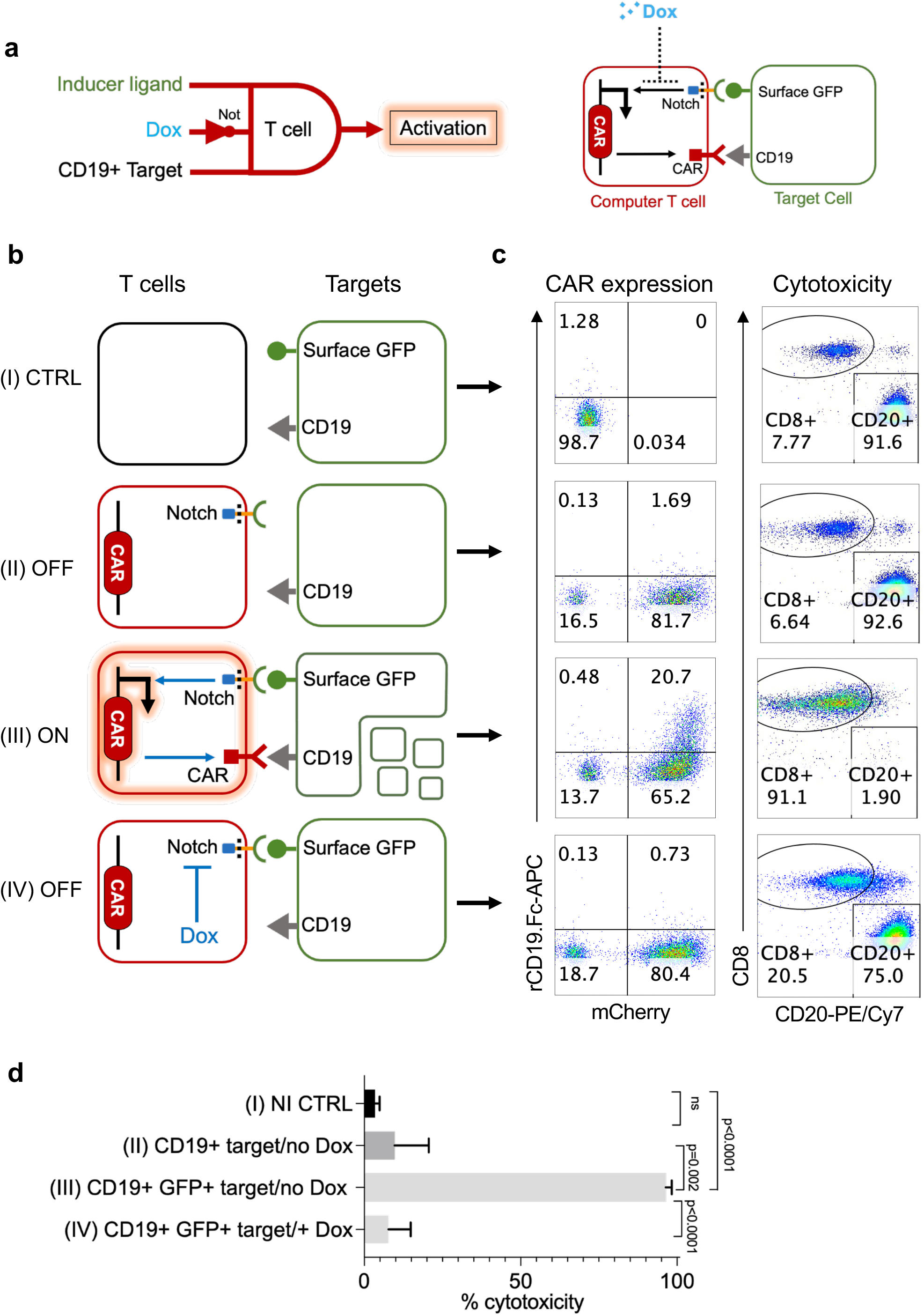
Integrating synthetic notch receptors into CAR-T failsafe toolbox. **a**, Illustration of engineered T cells becoming activated only after receiving the synthetic Notch (synNotch) ligand signal in the presence of CD19+ targets and where dox is absent (left) and their dox-repressible interaction with target cells that express the inducer Notch ligand (Surface GFP) and CD19 CAR antigen (right). The synNotch receptor expressed in engineered T cells consists of an anti-GFP extracellular domain that can bind to surface GFP on the inducer cells and an intracellular tetracycline-controlled transactivator (tTA) domain that would be released into the cytoplasm and transport into the nucleus to activate the TRE promoter upon ligand binding. The inducible CAR construct is under a TRE promoter and constitutively expresses mCherry as a marker for transduction. **b**, Cytotoxicity assay performed to evaluate the efficiency of synNotch CAR CD8+ T cells against T2 cells that express surface GFP ligand (left). The conditions include wild-type primary T cells against GFP+ T2 cells (I), synNotch CAR T cells against wild-type T2 cells (II), synNotch CAR T cells against GFP+ T2 cells in the absence (III) and presence (IV) of dox. **c**, SynNotch CAR CD8+ T cells would exhibit cytotoxicity only in the presence of inducer ligand and CD19 target antigen when dox is absent in the environment. The mCherry expression was used as a proxy to identify engineered T cells transduced with the TRE-CAR construct. Effector and target cells were identified with CD8 and CD20 surface expression, respectively. **d**, Comparing the killing by effector cells in the conditions mentioned in **b** after 72h (n=3). Cytotoxicity was calculated using the percentage of CD20 cells in the condition where wild-type T cells were combined with wild-type T2 cells. The error bars represent mean and one standard deviation. The experiments were replicated twice at different time points (n=3). An unpaired t-test was used to determine the significance in **d**.

We then explored the possibility of eliciting CD4+ T cell help to CD8+ T cells. For this purpose, we transduced primary human CD4+ T cells from the same donor with lentiviruses encoding the GFP ligand to make them “synNotch sender cells,” meaning the inducer cells of CAR expression in neighboring synNotch-CAR CD8+ T cells. We then set up a cytotoxicity assay, using wild-type CD4+ T cells and synNotch-CAR CD8+ T cells as controls (I), by co-culturing engineered GFP+ CD4+ T cells with synNotch-CAR CD8+ T cells and wild-type T2 cells in the presence (II) or absence (III) of dox for 3 days (Fig. 5a). synNotch-CAR CD8+ T cells only exhibited cytotoxicity when GFP+ CD4+ cells were present, and dox was absent (Fig. 5b, c, panel II). Taken together, these results showed that the synNotch system could be used to 1) activate CAR-T cells in response to a specific ligand, 2) regulate CAR-T cells with dox, and 3) enable intercellular cooperation of modified T cells against a common target.

**Figure 5.**
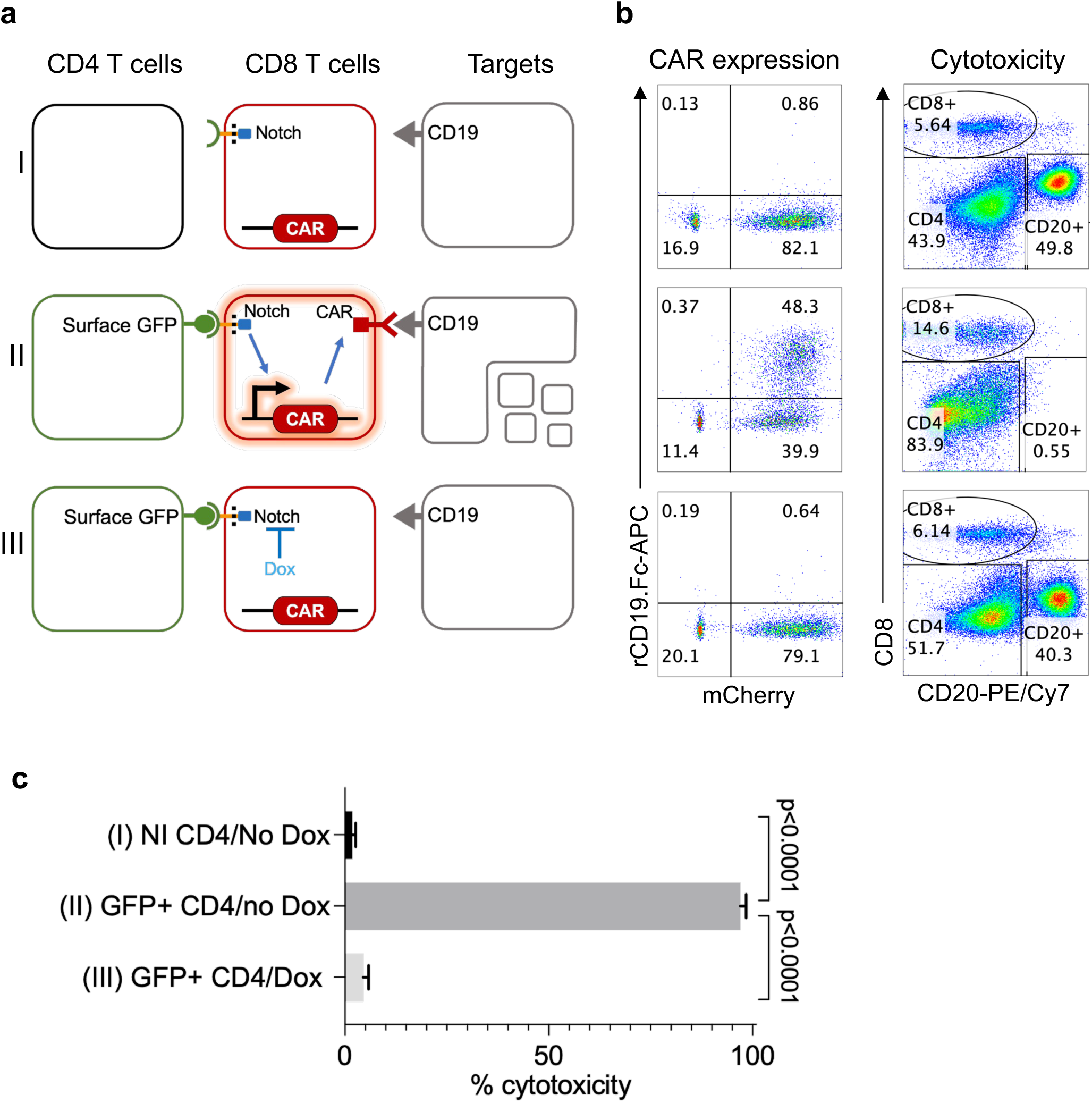
Triple cell-inducible notch system as a CAR-T cell fail-safe system. **a**, Schematic illustration of a cytotoxicity assay in which engineered CD4+ T cells were used to induce synNotch CAR T cells to express CD19 CAR which in turn would target T2 cells in the absence of dox. The conditions include wild-type primary CD4+ T cells used as a control combined with synNotch CAR T cells and CD19+ targets (i), GFP-expressing CD4+ primary T cells used as the inducer of synNotch CAR CD8+ T cells against CD19+ targets (ii) and GFP-expressing CD4+ primary T cells used as the inducer of synNotch CAR CD8+ T cells against CD19+ targets in the presence of dox (iii). The mCherry expression was used as a proxy to identify engineered T cells transduced with the TRE-CAR. **b**, Corresponding representative flow cytometry data showing the CAR expression in synNotch CAR T cells and the populations of CD4+, CD8+, and T2 cells. Effector and target cells were identified with CD8 and CD20 surface expression, respectively. **c**, Comparing the cytotoxicity percentages of effector cells as in Figure 4d after 72h Cytotoxicity was calculated using the percentage of CD20 cells in the condition where wild-type T cells were combined with wild-type T2 cells. The error bars represent mean and one standard deviation. The experiments were replicated twice at different time points (n=3). An unpaired t-test was used to determine the significance in **c**.

We next aimed to delete the CAR gene inserted in the genome of T cells using the CRISPR/Cas9 system as a permanent fail-safe measure to downregulate CAR expression. We first tested deleting CAR expression in a transformed CD4+ T cell line, Jurkat cells, to determine the best gRNA sequence for this goal. For this, five different guide RNA (gRNA) sequences targeting the CAR construct were designed (Supplementary Fig. 1a) and Jurkat cells engineered to stably express anti-CD19 CAR constructs were super-infected with CRISPR/Cas9 lentiviruses constitutively encoding Cas9 and engineered gRNAs. A CRISPR construct carrying an empty gRNA vector was used as a control. Six days following the CRISPR lentivirus transduction, the cells were collected and stained with antibodies for their CAR expression. While all the gRNAs reduced the CAR expression, C2 and C4 clones eliminated nearly all CAR expression in Jurkat cells (Supplementary Fig. 1b). The C2 gRNA was subcloned into a dox-inducible tet-on CRISPR/Cas9 expression vector and transduced into primary human CD8+ T cells with the CAR construct. The cells were then cultured for 10 days and stained for their CAR expression. The CAR-expressing CD8+ T cells transduced with lentivirus encoding dox-inducible CRISPR showed a substantial reduction in CAR expression even without dox, suggesting a significant level of leakiness of the promoter (Supplementary Fig. 1d, panel II) Treatment with 1 ug/mL of dox for nine days significantly reduced CAR expression compared to controls (Supplementary Fig. 1d, e, panel IV). However, T cells continued to lose significant levels of CAR expression after nine days in culture compared to initial expression (Supplementary Fig. 1d, panel III). While we tried various ways to express these Cas9 and gRNA constructs (including in separate vectors, data not shown), we could not achieve better results, suggesting this approach may not be a viable fail-safe switch.

Lastly, we explored a potential killer switch for CAR-T cells with a novel concept by creating fratricide among CAR-T cells, which we named the “battle royale” approach. For this, instead of engineering a suicide gene into the CAR-T cells, we engineered controllable expression of surface CD19 antigen in the same cells that also express the anti-CD19 CARs. Thus, the induction of CD19 on CAR-T cells would render them targets of other neighboring anti-CD19 CAR-T cells (Fig. 6a). Accordingly, human CD8+ T cells were co-infected with lentiviruses encoding a constitutively expressed CAR and doxycycline-inducible CD19 constructs and then cultured for 14 days. Primary CD8+ T cells engineered with lentiviruses encoding a non-functional CAR with deleted intracellular signaling domains (deadCAR, dCAR, Supplementary Fig. 2a) were used as controls. The cells were then treated with 0 or 3 ug/mL of dox for two days and stained with anti-CD19 to determine the level of CD19 induction in T cells (Fig. 6b). In addition, cells were stained with anti-CD25 and with rCD19.Fc antigen to determine the levels of T cell activation and expression of CAR on these cells, respectively (Fig. 6c). Upon dox treatment, all conditions upregulated CD19 expression (Fig. 6b). However, as expected, there was much higher CD19 expression in both no-CAR and dead-CAR T conditions compared to functional CAR cells, as the latter cells would be expected to kill each other that expressed CD19 (Fig. 6b, top panel). Consistent with this, CAR-T cells with the inducible CD19 construct were also robustly activated based on CD25 upregulation, whereas dead-CAR and no-CAR conditions did not display any T cell activation (Fig. 6c).

**Figure 6.**
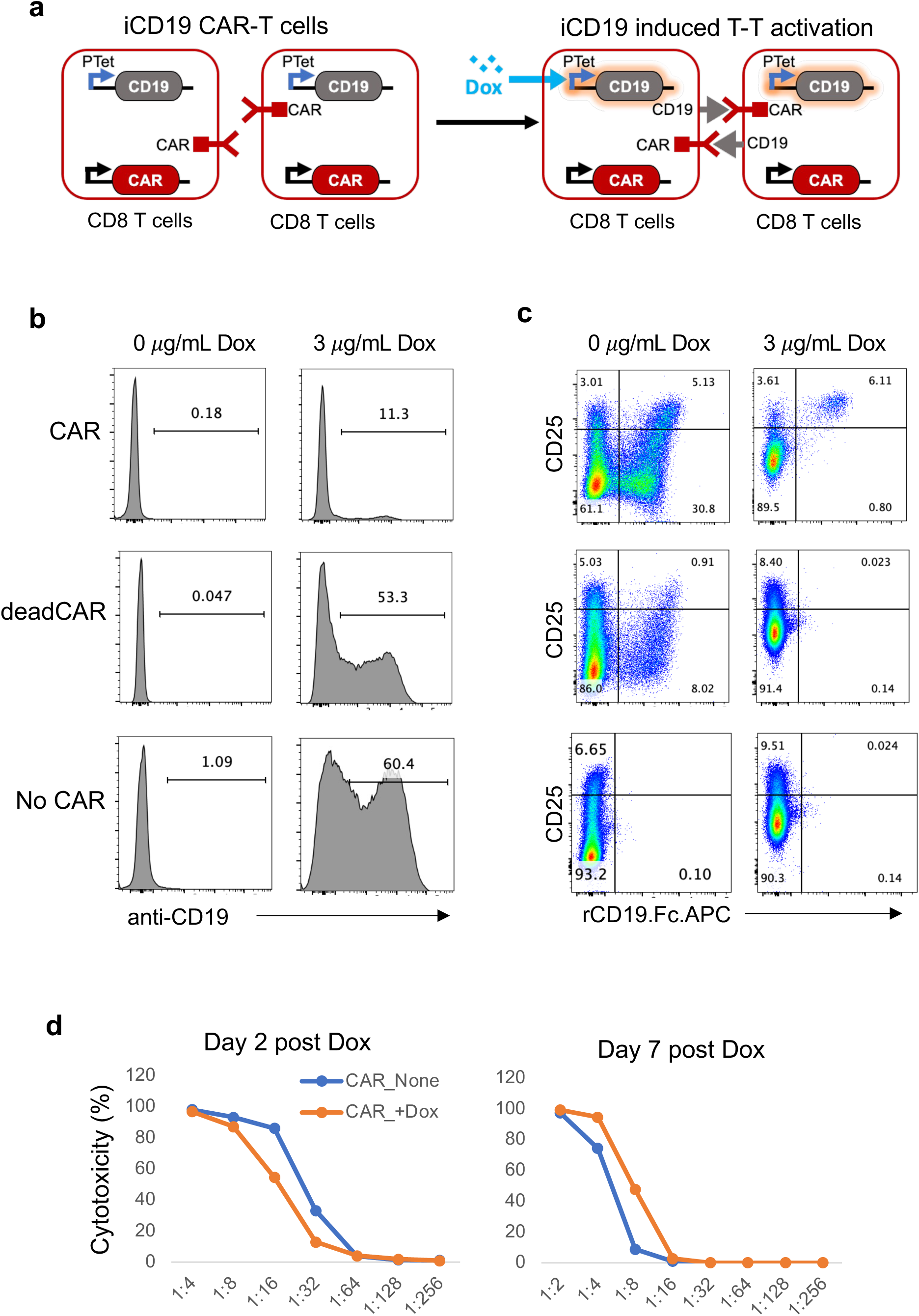
Fratricide ‘Battle Royale” system for CAR-T cells. **a**, Schematic representation of inducible CAR-T cell elimination system. Engineered T cells equipped with CD19-CAR and dox-inducible CD19 constructs would target each other and reduce the number of CAR-T cells in the presence of dox. **b**, Cytotoxicity assay in which CAR-T cells were induced to express CD19 on their surface for “battle royale” fratricide (left) and the corresponding representative flow cytometry data showing the CD25 and CAR expressions of the engineered T cells in the absence and presence (3 *μ*g/ml) of dox (right). The experiment conditions include CAR-T cells with inducible CD19 construct (top panel), engineered T cells with inducible CD19 construct and a CAR construct that has no intracellular domain (dead CAR, dCAR) (middle panel), and engineered T cells with empty CAR vector and inducible CD19 construct (bottom panel). Surface CAR, CD19 and CD25 expressions were used to evaluate the cytotoxicity and the activation of the cells. **c**, Flow cytometry plots showing the CD25 and CAR expressions of the conditions mentioned in **b** in the presence (3 *μ*g/mL) and absence of dox (n=3)**. d**, Line graphs demonstrating the percent cytotoxicity of the non-induced (blue), in the presence of doxycycline, and non-induced (orange), in the absence of doxycycline, CAR-T cells (mentioned in the top panel in c) against CD19+ targets. X axis shows the effector to target ratio (E:T) of the cytotoxicity assay (n=3). The experiments were replicated twice at different time points (n=3).

We next determined the cytotoxic potential of the CD19-induced CAR-T cells 2 and 7 days after dox induction (Fig. 6d). As expected, 2 days after dox induction, the cytotoxicity of induced cells was significantly lower compared to non-induced parents (Fig. 6d). However, after seven days, CD19-induced CAR-T cultures displayed higher cytotoxicity at different effector-to-target ratios (Fig. 6d). We reasoned that because during the “battle royale,” surviving CAR-T cells were activated, they could also be proliferating and increasing in proportions. Indeed, upon staining the cells for CAR expression, the proportion of CAR-expressing cells was greatly reduced after one day in dox but rapidly increased to recover to original levels by day seven and further continued to increase by day 14 post dox-induction (Supplementary Fig. 2b).

It was, however, still puzzling that the cytotoxicity of induced CAR-T cells at day 2 was not reduced relative to their proportion in the culture and, conversely, at day 7 showed higher cytotoxicity despite being in similar proportions to uninduced cultures (Fig. 6c and Supplementary Figure Fig. 2b). One possibility for this difference could be that induced CAR-T cells, being activated, could be more robust in displaying their cytotoxic activity upon reactivation with target cells. Alternatively, the surviving cells could be less exhausted and thus more potent in their functional responses. Given the difficulty of determining this in mixed bulk culture experiments, we tested this hypothesis utilizing a novel approach using the Enrich TroVo system^28^. This approach is predicated on the competitive selection of both surviving/proliferating and cytotoxic CAR-T cells from a mixed population.

To evaluate the functional disparities among CAR-T cells, such as their tumor cytolytic activity and proliferative potential, we utilized a repeated antigen exposure assay^29^, which involve repeated tumor rechallenge every three days. The Enrich TroVo system facilitated the creation and monitoring of approximately 500 micro hydrogel chambers to conduct these repeated tumor exposure cocultures (Fig. 7a). Through marker-free imaging and the utilization of anti-CD8 staining of T cells, we were able to identify tumor cells and CAR-T cells, respectively, as shown by red spots (Fig. 7a). The confluence of T cells and tumor cells within these cultures was meticulously monitored for 10 days. The experimental design included both uninduced and induced CAR-T cells, alongside induced deadCAR and No CAR controls (Fig. 7a).

**Figure 7.**
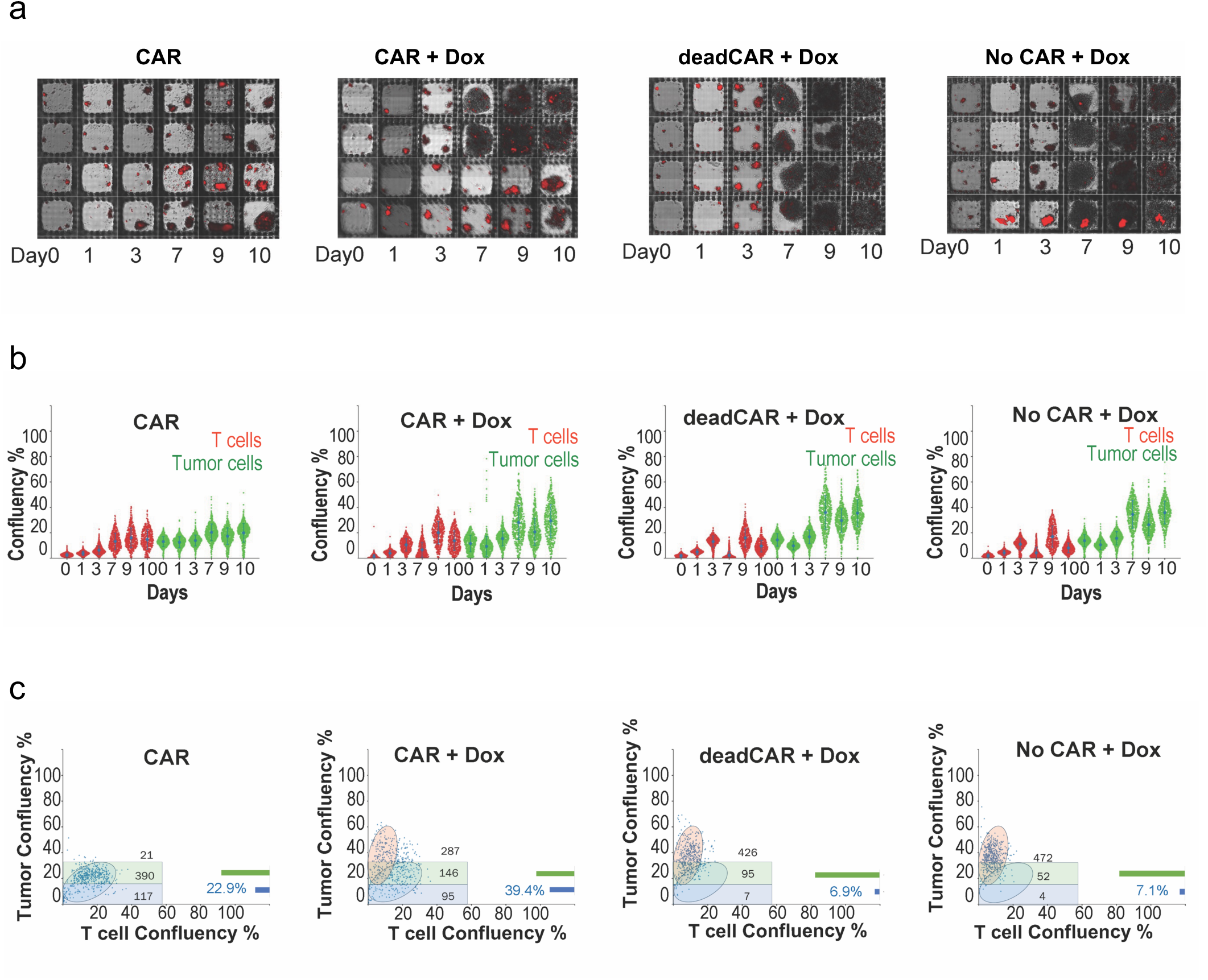
Monitoring CAR-T cell ‘Battle royale” using Enrich Trovo system. **a,** CAR-T cell chronic antigen exposure assay using Enrich TROVO system where cell movement is restricted (3) to small individual “rooms” (250μmx250μm) made by biocompatible hydrogels. For each room, 5-10 T cells were cocultured with 15-30 tumor cells initially and ∼10 tumor cells were fed to each room every 3 days to evaluate persistent T cell function. TROVO built-in imaging and quantification software are used to generate a movie. **b,** T cell (anti-CD8 antibody-AXE647)/Tumor cell (marker free detection) quantification for each room in sina plots. To simultaneously monitor tumor suppression and T cell proliferation, T cell and tumor cell amounts were measured by the confluency as y axis on day 0, 1, 3 (tumor feed 1), 7 (tumor feed 2), 9 (Tumor feed 3), 10 of uninduced (CAR) and induced (battle royale, CAR + Dox), two negative controls (deadCAR + Dox, No-CAR + Dox) and visualized by scatter-plot**. c,** Dot graphs in which the percent tumor and T cell confluency values at day 7 were plotted together. T cells were separated in to 3 different groups. highest tumor confluency. Based on this data, T cells were divided into three parts: 1. Top-performing CAR-T cells (blue shade), 2. Medium-performing CAR-T cells (green shade), and 3. non-killers (pink shade). Percentage values shown correspond to the top performing CAR-T cell frequency in their respective microwells at day 7. The experiments were replicated twice at different time points (n=3).

Our observations revealed that a significant majority (>90%) of the uninduced CAR-T cells effectively suppressed tumor cell proliferation, whereas the negative controls exhibited substantial tumor growth, indicative of minimal cytotoxicity (Fig. 7b). Following two rounds of tumor rechallenge (Day 7), the induced CAR-T cells segregated into two distinct phenotypes based on their tumor-killing capabilities by day 10 (Supplementary Fig. 3). Upon further tumor rechallenge (Day 7), we reanalyzed this data by dividing the T cell and tumor confluency into three parts: 1. Top-performing CAR-T cells (Fig. 7c, blue shade), 2. Medium-performing CAR-T cells (Fig. 7c, green shade), and 3. Non-killers (Fig. 7c, pink shade).

We found that wells containing top-performing CAR-T cells (Fig. 7c, blue shades), characterized by the suppression of tumor confluence to below 16% (threshold determined from the negative control), were higher (39%) compared to non-induced CAR-T wells (23%), while the frequency in controls was about 7% (Fig. 7c). As expected, about 50% of the wells in the induced CAR-T condition did not show tumor killing, given that cells in these T cells had completely died during the fratricide. Together, these experiments underscore the potential of the “battle royale” process as a viable strategy for enhancing the selection and enrichment of potent CAR-T cells within a heterogeneous population.

## Discussion

In this study, we developed a synthetic biology toolbox for engineering CAR-T cells with tunable and controllable features. We combined different genetic circuits to achieve temporal regulation of CAR expression and activity using dox as a common modulator. We also introduced intercellular communication and collaboration among modified T cells using synthetic notch receptors and engineered CD4+ T cells to help cytotoxic CD8+ T cells. Furthermore, we developed a fratricide-based approach both initially reducing and subsequently expanding for more potent CAR-T cells.

Our toolbox provides diverse solutions for improving the safety and efficacy of CAR-T cell therapies by integrating different genetic circuits with remote modulators. First, we showed that dox can induce or repress CAR expression in iOnCAR or iOffCAR approaches, respectively, allowing for flexible adjustment of CAR-T cell function in clinical settings depending on the tumor burden and CAR-T cell toxicity profile. This is consistent with previous studies that have used Tet systems to regulate CAR expression or activity in response to dox^18,20,30,31^. Importantly, our iOnCAR T cells were highly competent at killing target cells even when only 20% of them were expressing CAR on their surface and did not show cytotoxicity in the absence of dox, suggesting that iOnCAR could be a tightly controlled system for CAR-T regulation.

We have also shown that dox can modulate synNotch receptor-mediated CAR induction by an antigen on the target cells, enabling fine-tuning of antigen-specific activation and reducing on-target off-tumor toxicity risk. By coupling synNotch receptors with CARs, we have achieved antigen-specific activation of CAR-T cells, only when the cells express both antigens. This study corroborated reports by others that used synNotch receptors to induce CAR expression^23,27,32^; however, our toolbox is novel in its use of dox to remotely modulate synNotch receptor activity. This allows for more precise control over CAR-T cell function and potential fine-tuning. It may also reduce the risk of on-target off-tumor toxicity by avoiding bystander tissues that share the CAR-T target antigen via local dox administration. In a reverse strategy, using rtTA instead of tTA in the Notch receptor to turn on the system in the presence of dox, CAR-T cells could be activated only in the tumor microenvironment via local dox administration. These approaches would enable both spatial and temporal regulation of CAR-T cells in vivo.

Our novel approach to express a synthetic ligand (GFP) in CD4+ T cells to induce CAR expression in synNotchCAR CD8+ T cells was aimed at mimicking natural cooperation between helper and cytotoxic T cells, under the control of dox. We showed that synNotchCAR CD8+ T cells only exhibit cytotoxicity when GFP+ CD4+ cells are present and dox is absent. This enables intercellular cooperation of modified T cells against a common target and provides an additional layer of regulation for CAR-T cell function. Moreover, this approach can potentially enhance the efficacy and persistence of CAR-T cell therapies by exploiting the helper function of CD4+ T cells, which can provide cytokines and costimulatory signals to CD8+ T cells in the tumor microenvironment^33–36^. Further, this approach may stimulate the utilization of engineered CD4+ T cells for cancer immunotherapy, in addition to cytotoxic CD8+ T cells. It is also possible that the surface GFP molecule which CD4+ T cells are using to induce CAR expression on CD8+ T cells could be engineered with another synthetic Notch receptor that recognizes a second tumor marker, as a further fail-safe mechanism. Further studies will be needed to explore the possible utility and safety of these methods in *in vivo* models.

We also tried a CRISPR-mediated approach for the permanent elimination of CAR expression should they show high toxicity. While we could show that dox can induce the depletion of CAR construct in primary T cells via CRISPR/Cas9. We also observed some background activity on the inducible CRISPR system that reduced CAR expression even without dox treatment, suggesting continuous level of leakiness in Cas9 expression. To be useful as a viable fail-safe approach, this approach will require using alternative methods to regulate CRISPR/Cas9 expression.

Finally, we have evaluated a novel approach, referred as “battle royale”, that involves inducing fratricide within engineered CAR-T cells, as a means of reducing their numbers, should the need arise in clinical settings. We have shown that dox can induce CD19 expression in CAR-T cells, which in turn triggers cytotoxicity by other CAR-T cells. A potential advantage of this approach is that it would work preferentially in cancer tissue where CAR-T cells are within the same milieu, thus sparing other CAR-T cells because this system requires at least two CAR-T cells to be neighboring each other. In addition, T cell-mediated cell death, as opposed to suicide switches, is a naturally occurring phenomenon that is activated when T cells recognize each other as foreign or infected ^37,38^, and may produce less inflammatory signaling than mass suicide induction *in vivo*. Another important aspect of the “battle royale” approach is that, while in the short term it purges a significant portion of the CAR-T cells through fratricide, the surviving and activated CAR-T cells rapidly expand and reach similar numbers after about a week of induction, continuing to increase for at least another week in in vitro cultures. Remarkably, upon this activation and expansion, these cells also display higher cytotoxicity to target cells on an individual cell basis, as demonstrated using the Enrich TroVo system. It is conceivable that during the initial battle royale killing, those CAR-T cells with an exhausted phenotype are more likely to be eliminated, resulting in the expansion of cells that are more robust in their functions upon reactivation with the targets. Thus, it is tempting to speculate that the battle royale approach can be used to enhance in vivo CAR-T cell responses for the long term and evade T cell exhaustion.

Taken together, we have developed a multi-layered fail-safe toolbox for programming CAR-T cells with tunable and controllable features. Our toolbox provides diverse solutions for improving the safety of CAR-T cell therapies by integrating different genetic circuits with remote controllers. These synthetically programmed T cells facilitate intercellular communication and collaboration, introducing novel means for regulating these potent cells and utilizing non-cytotoxic T cells to aid the targeting of malignancies by cytotoxic T cells. The fail-safe toolbox we developed in this study, leveraging the advantages of synthetic biology, could potentially be applied to other cell-based immunotherapies or gene therapies that require precise and adaptable control over gene expressions and various cell functions.

## Materials and Methods

### CAR cassette

The CAR construct used in this study consists of; CD8 alpha signal peptide, a single chain variable fragment (scFv) of anti-CD19 or anti-Spike protein antibodies, CD8 hinge domain, CD8 transmembrane domain, 4-1BB (CD137) intracellular domain and CD3ζ domain. It was designed with Snapgene and synthesized (Genscript). The CD8a signal peptide, CD8 hinge, CD8 transmembrane domain, 4-1BB intracellular domain and CD3ζ domain sequences were obtained from Ensembl Gene Browser and codon-optimized with SnapGene by removing the restriction enzyme recognition sites that are necessary for subsequent molecular cloning steps, while preserving the amino acid sequences. Anti-CD19 scFv amino acid sequence was obtained from Addgene plasmid #79125, reverse translated to DNA sequences and codon-optimized with Snapgene 5.2.4. The constructs were then cloned into respective lentiviral expression vectors using relevant restriction enzymes.

### iOnCAR, iOffCAR, TRE-CAR and dead CAR Constructs

iOnEmpty vector was built into the TLCV2 plasmid from Addgene (#87360) by amplifying and cloning its own tight TRE promoter into the backbone and removing the U6 Promoter-empty gRNA-Tight TRE Promoter-Cas9 cassette via KpnI-BamHI restriction digestion. To make iOnCAR, the CD19-CAR cassette was then cloned into iOnEmpty vector using EcoRI-BamHI restriction sites while keeping the T2A-GFP downstream peptides in-frame to preserve the reporter gene co-expression. The iOffEmpty, and subsequently the iOffCAR constructs were built on iOnEmpty by swapping the rtTA with tTA from the pHR_PGK_antiCD19_synNotch_TetRVP64 vector from Addgene (#79126) via XbaI-XhoI sites. TRE CAR construct was generated from pHR_Gal4UAS_pGK_mCherry (Addgene #79124) by inserting a TRE promoter and removing GAL4 UAS promoter via EcoRI restriction digestion and cloning CD19 CAR construct into the multiple cloning site. Dead CAR cassette was generated via amplifying only the signal peptide, extracellular and transmembrane domains of CD19 CAR construct and adding a stop codon after transmembrane with PCR. The PCR product was then cloned into the lentivector backbone to be used as a control to the CD19 CAR lentivector.

### CRISPR constructs

The sequence of gRNAs used for gene knockout were designed using the IDT CRISPR tool (https://www.idtdna.com/site/order/designtool/index/CRISPR_CUSTOM). All sequences were selected to precede 5′-NGG protospacer-adjacent motif (PAM) sequence. Cloning of gRNAs into LentiCRISPR v2 was modified from Sanjana *et al* ^39^. The forward and reverse primers used in the generation of double-strand, sticky-ended gRNA inserts were designed using an open-sourced web tool automating the original protocol (https://jsfiddle.net/mdmikaildogan/umbyux60/1/). The lentiviral CRISPR plasmids were digested with *BsmBI* (Thermo Fisher Scientific) for 30 min at 37°C. The digested plasmids were gel purified using Agarose Gel and DNA Gel Extraction kit (Monarch), according to the manufacturer’s recommendations. The forward and reverse oligonucleotides that encode the gRNAs (Eurofins Genomics) were annealed and phosphorylated in a mixture of T4 Ligation buffer and T4 PNK at 37°C for 30 min followed by heat inactivation at 95°C for 5 min, then ramp down to 25°C at 5°C/min. Diluted annealed oligos were ligated to digested plasmids in Ligase Buffer at room temperature for 10 min. The cloned constructs were then transformed into NEB® Stable Competent *E. coli* (High Efficiency) (New England Biolabs) according to the manufacturer’s protocol. The colonies were cultured overnight for plasmid DNA isolation using QIAprep Spin Miniprep Kit and Qiacube (Qiagen). Diagnostic digest was performed for confirming the positive clones: the purified colonies with LentiCRISPR v2 and TLCV2 backbones were digested with both EcoR1 and BamHI restriction enzymes. The colonies with positive insertion were confirmed by analyzing the resulting fragments by gel electrophoresis.

### Lentivirus production

The lentiviruses pseudotyped with vesicular stomatitis virus G protein envelope were generated with HEK293T cells. Briefly, the lentivector plasmids containing the constructs were co-transfected with vesicular stomatitis virus G protein, pLP1, and pLP2 plasmids into HEK293T cells at 80–90% confluency using Lipofectamine 3000 (Invitrogen) according to the manufacturer’s protocol. The transfection medium was replaced with RPMI 1640 with 10% Fetal Bovine Serum (FBS, Atlanta Biologicals) 6 hours post-transfection. Viral supernatants were collected 48 hours post-transfection and filtered through a 0.45-μm syringe filter (Millipore) to remove cellular debris. A Lenti-X concentrator (Takara Bio USA) was used according to the manufacturer’s protocol to concentrate the virus 10-20X and the resulting lentiviral stocks were aliquoted and stored at −80°C. To measure viral titers, the virus preparations were serially diluted on Jurkat cells and 3 days post-infection, infected cells were measured using flow cytometry and the number of cells transduced with 1 ml of virus supernatant was calculated as infectious units/ml. 72 hours after infection, GFP-positive cells were counted using flow cytometry and the number of cells transduced with virus supernatant was calculated as infectious units/ml. Based on these titer values, primary T cells, 293 T cells and T2 cells were transduced with a multiplicity of infection (MOI) of 3-10.

### Engineering CAR-T cells and Surface GFP-expressing target cells

Healthy adult blood was obtained from AllCells. PBMCs were isolated using Ficoll-paque plus (GE Health care). CD8+ T cells were purified using Dynal CD8 Positive Isolation Kit (from Invitrogen). CD8+ T cells were >99% pure and assessed by flow cytometry staining with CD8-Pacific Blue antibody (Biolegend). Total CD8+ T cells were activated using anti-CD3/CD28 coated beads (Invitrogen) at a 1:2 ratio (beads:cells) and infected with their respective lentiviral constructs with a multiplicity of infection (MOI) of 5-10. The cells were then expanded in complete RPMI 1640 medium supplemented with 10% FBS, 1% penicillin/streptomycin (Corning Cellgro) and 20ng/ml of IL-2, cultured at 37°C in 5% CO_2_ supplemented incubators. Respective viruses were added 24 hours after the activation. Cells were expanded for 10-12 days and cytotoxicity assays were performed following their expansion. All engineered and wild-type T2 cells were cultured in complete RPMI 1640 medium supplemented with 10% FBS, 8% GlutaMAX (Life Technologies), 8% sodium pyruvate, 8% MEM vitamins, 8% MEM nonessential amino acid, and 1% penicillin/streptomycin (all from Corning Cellgro). To generate T2 cells with stable surface GFP overexpression, wild-type T2 cells were transduced with the surface GFP overexpressing lentivirus at an MOI of 3 and proliferated. The infection levels were determined by GFP expression using flow cytometry analysis.

### Staining and flow cytometry analysis

Cells were resuspended in staining buffer (PBS + 2% FBS) and incubated with fluorochrome-conjugated antibodies for 30 min at 4°C. CD8+ T cells were identified with CD3-Pacific Blue (cat# 300431) or CD8-Pacific Blue (cat# 301033) antibodies (Biolegend). Activation of CD8+ CAR-T cells was determined with CD25 staining using CD25-APC antibody (Biolegend, cat# 302610). CAR expression of anti-CD19 CAR was determined with Human CD19 (20-291) Protein, Fc Tag, low endotoxin (Super affinity) (Acro) followed by secondary staining with APC conjugated anti-human IgG Fc Antibody (Biolegend, cat# 410712) and GFP/RFP expressions. For cytotoxicity assay analysis, stably surface GFP-expressing T2 cells were identified with the GFP marker or with their CD19 and CD20 expressions via PE/Cy7-CD19 (Biolegend, cat# 302216) and PE/Cy7-CD20 (Biolegend, cat# 302312) antibodies, respectively. Samples were acquired on a BD FACSymphony A5 analyzer and data were analyzed using FlowJo (BD Biosciences).

### Cytotoxicity assay

Following the expansion of engineered CAR-T cells for 10-12 days, the cells were analyzed for their GFP and CAR expressions. Effector-to-target (E:T) cell ratio was calculated based on the number of CAR-expressing cells. The CAR-expressing cells were titrated from 2:1 to 1:16 effector to target cell ratio at 2-fold dilutions while the target cell number was constant. Cytotoxicity assay conditions were analyzed with flow cytometry at 72 hours post-coculture and the cells were identified as described in Staining and flow cytometry analysis. Cytotoxicity for each E:T condition was calculated based on the corresponding control condition with the following formula: (Percent of target cells in control condition – percent of target cells in experimental condition) / percent of target cells in control condition * 100.

### Enrich Trovo system

#### Microwell Printing

In a 24-well glass bottom plate, around 500 hydrogel microwells (250 um x 250 um) were printed (MatTek) using 100 uL of prewarmed Microwell Maker (Enrich Biosystems) per well. Microwells were printed using Trovo with the following optimized parameters: 1.9 s exposure, 13 mm diameter, and 2.0 step size. After printing, each well was washed with 1 mL of 1X Phosphate Buffer Saline (PBS) (Sigma-Aldrich) and incubated at 37°C for 5 min. After incubation, PBS was aspirated from the wells. Washing steps were repeated two more times until residual gel was removed. The plate was then stored in the 37°C incubator until use.

#### Co-culture Setup

Uninduced iCD19-CAR-T cells with T2 Lymphoma were seeded in the microwells in their respective wells in a total of 50 uL of complete medium containing Alexa Fluor 647 CD8a Antibody (RPA-T8, Thermo Fisher) per well. Effector to target (E:T) ratios of 1:2 was used on average of 20 T cells per micro well. After one hour of cell settling and T-cell staining, wells were prepared for imaging by washing with 300 uL complete medium.

#### Microwell Chronic Antigen assay

Co-cultures were imaged with Trovo^TM^ (Enrich Biosystems) on Day 0, 1, 3, 7, 9 and 10. Each time the plate was imaged, T-cells were stained for one hour before with A647 CD8 antibodies, followed by washing. After imaging, wells were kept in 500 uL of complete medium with 100 U IL-2. On Day 0, co-cultures were imaged using the GFP channel on Trovo to confirm induction of T cells. Subsequently, fluorescent marker A647 and brightfield were used for the rest of the experiment. Every 3 days, target tumor cells were readded after imaging with the addition of on average about 10 T2 cells to each well.

#### Imaging and data analysis

For each dataset, composite images with a 6x magnification encompassing both brightfield and fluorescence modalities were acquired utilizing the automated scanning and focusing capabilities of the Enrich TroVo platform. Subsequently, the system’s automatic microwell analysis algorithm facilitated the identification, segmentation, and indexing of individual microwells. Moreover, microwell-based cinematic sequences were autonomously compiled upon the completion of multi-day imaging datasets. Quantitative assessments of tumor cell and T cell populations, including cell counts and confluence metrics, were executed and subsequently tabulated within a comma-separated values file. These data were further elucidated through temporal representation via Sino-plot time series and daily bidimensional scatter plots, serving to monitor T cell proliferation and tumor cell suppression kinetics, facilitated by the automated data analysis functionality of the Enrich TroVo system.

### Statistical Analyses and Reproducibility

All statistical analyses were performed, and graphs were prepared using GraphPad Prism V9 software. The central tendency (mean) and the variations (standard deviation and standard error of mean) were also calculated using GraphPad Prism V9 software. The numbers of repeats for each experiment were described in the associated figure legends.

## Acknowledgments

We thank Dr. Sara Cassidy for the critical reading of this manuscript.

## Data availability

The data that support the findings of this study are available from the corresponding author upon reasonable request.

## Inclusion and Ethics and statement

We affirm that this study adheres to the highest ethical standards, with all procedures performed in accordance with the relevant guidelines. Our team is committed to fostering diversity and inclusion in all aspects of our research. We have striven to minimize bias in our study design, implementation, and analysis and are transparent about our methodology.

---

**Supplementary Figure 1.**
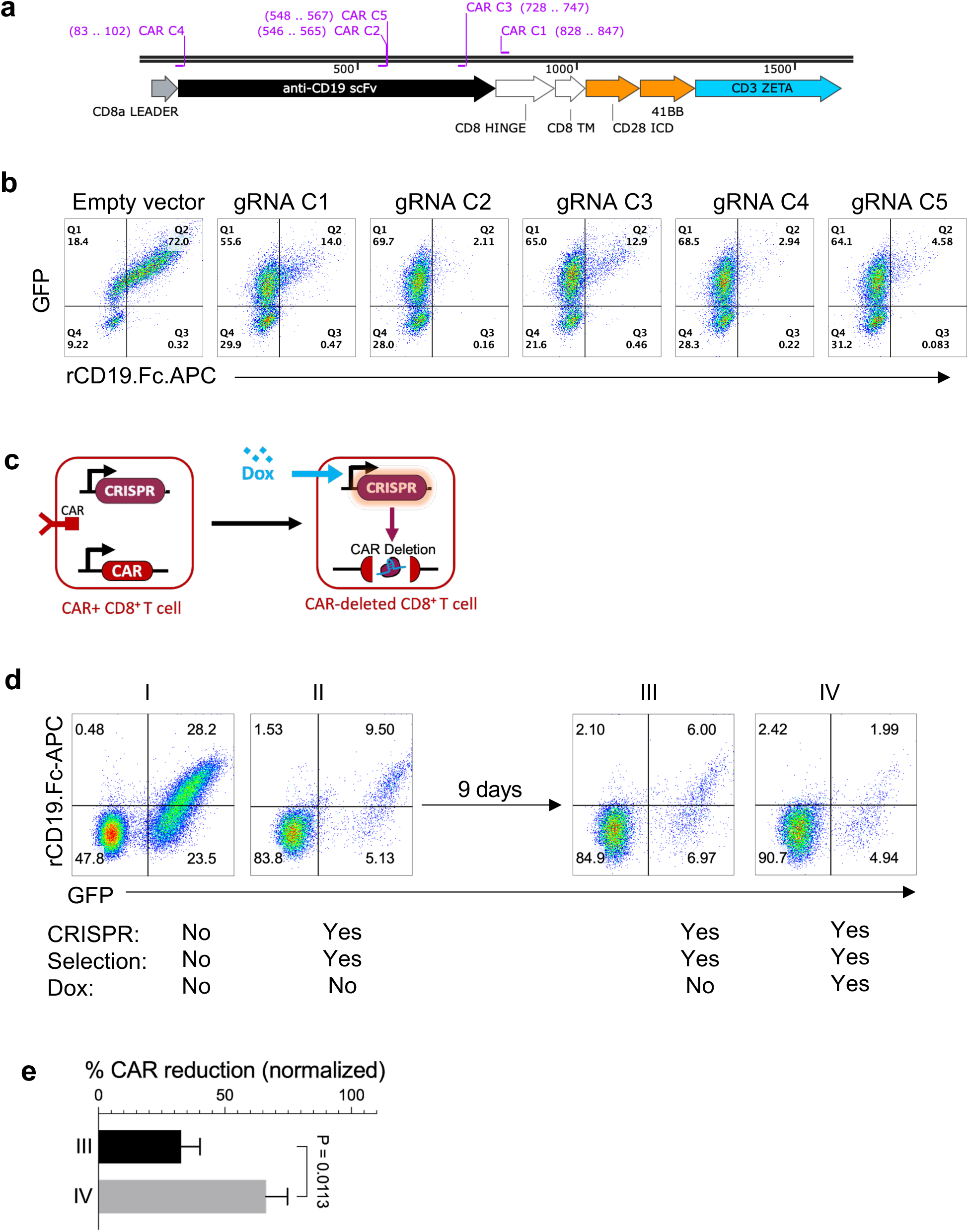
Development of CRISPR guide RNAs against CD19 CAR. **a**, CD19 CAR construct by which CRISPR gRNA targeting sites were shown. **b**, CRISPR-mediated depletion of CAR expression on Jurkat cell line to evaluate the functionality of the gRNAs. Empty CRISPR vector used as a control. Jurkat cells engineered to express the CD19 CAR construct were transduced with CRISPR vectors encoding Cas9 and custom gRNAs targeting different areas of the CAR construct. The cells were stained for their CAR expression as described in the methods six days after CRISPR transduction. **c,** Schematic illustration of CAR deletion via dox-inducible CRISPR/Cas9 system. Dox-inducible CRISPR construct consists of a gRNA targeting the CD19 CAR construct, a tet-on inducible Cas9, and constitutively expressed puromycin as a selection marker. **d**, CAR expression depletion after puromycin selection in the presence (1*μ*g/mL) or absence of dox. **e**, CAR expression in the presence versus in the absence of dox. Reduction in CAR expression was normalized using the control condition. The error bars represent mean and one standard error of mean. The experiments were replicated twice at different time points (n=3). A paired t-test was used to determine the statistical significance.

**Supplementary Figure 2.**
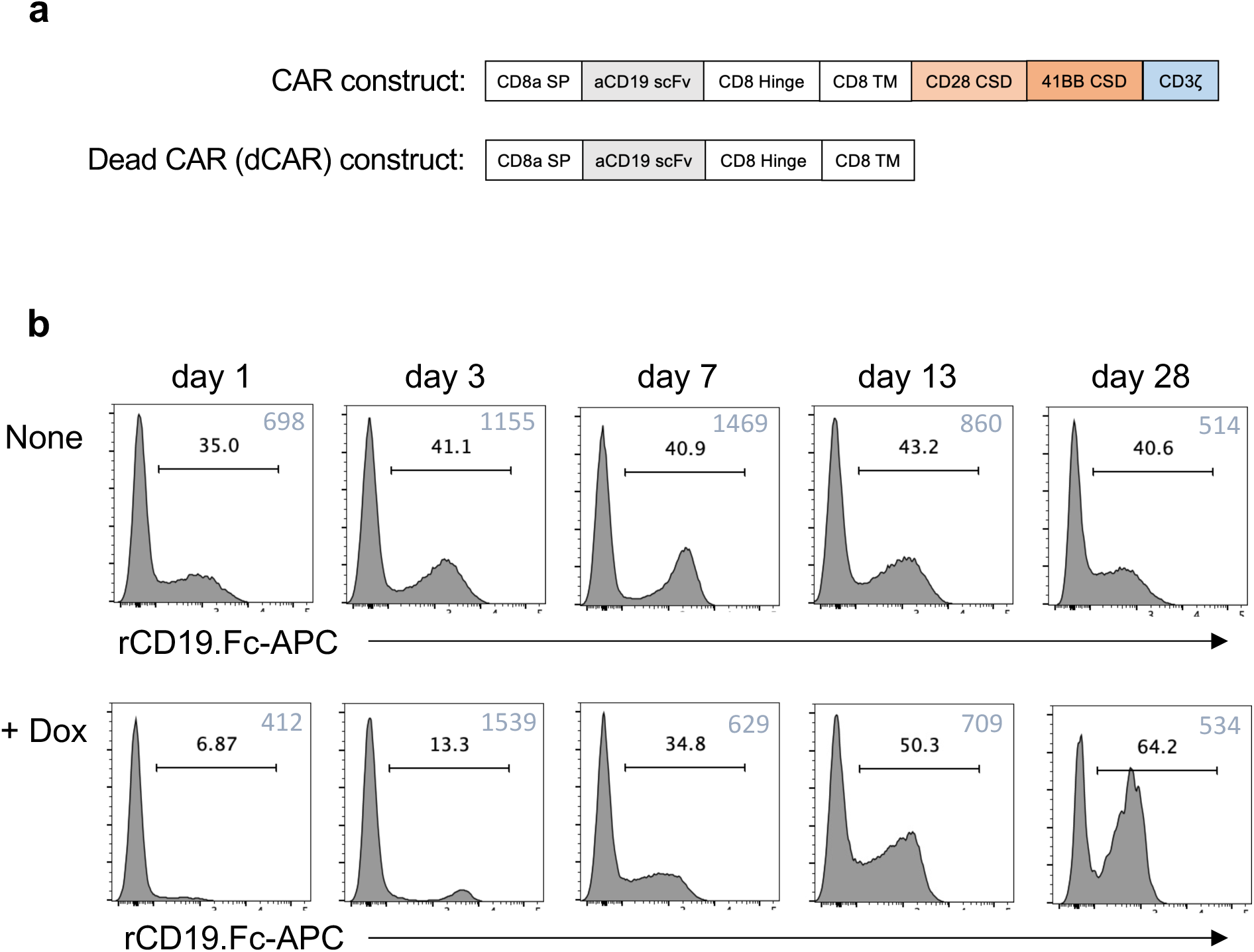
Map of a control CAR construct lacking intracellular costimulatory and activation domains. **a,** An outline of the dead CAR (dCAR) construct in comparison with CD19 CAR. dCAR construct lacks the intracellular domains of CD19 CAR and hence could be used as a control to CD19 CAR. **b,** Surface expression of CAR on T cells engineered with CAR and inducible CD19 constructs in the presence and absence of dox in different time points from day 1 to day 28 post doxycycline or control media addition. Cells were stained with recombinant CD19-Fc followed by anti-Fc APC, analysis was performed at indicated days.

**Supplementary Figure 3.**
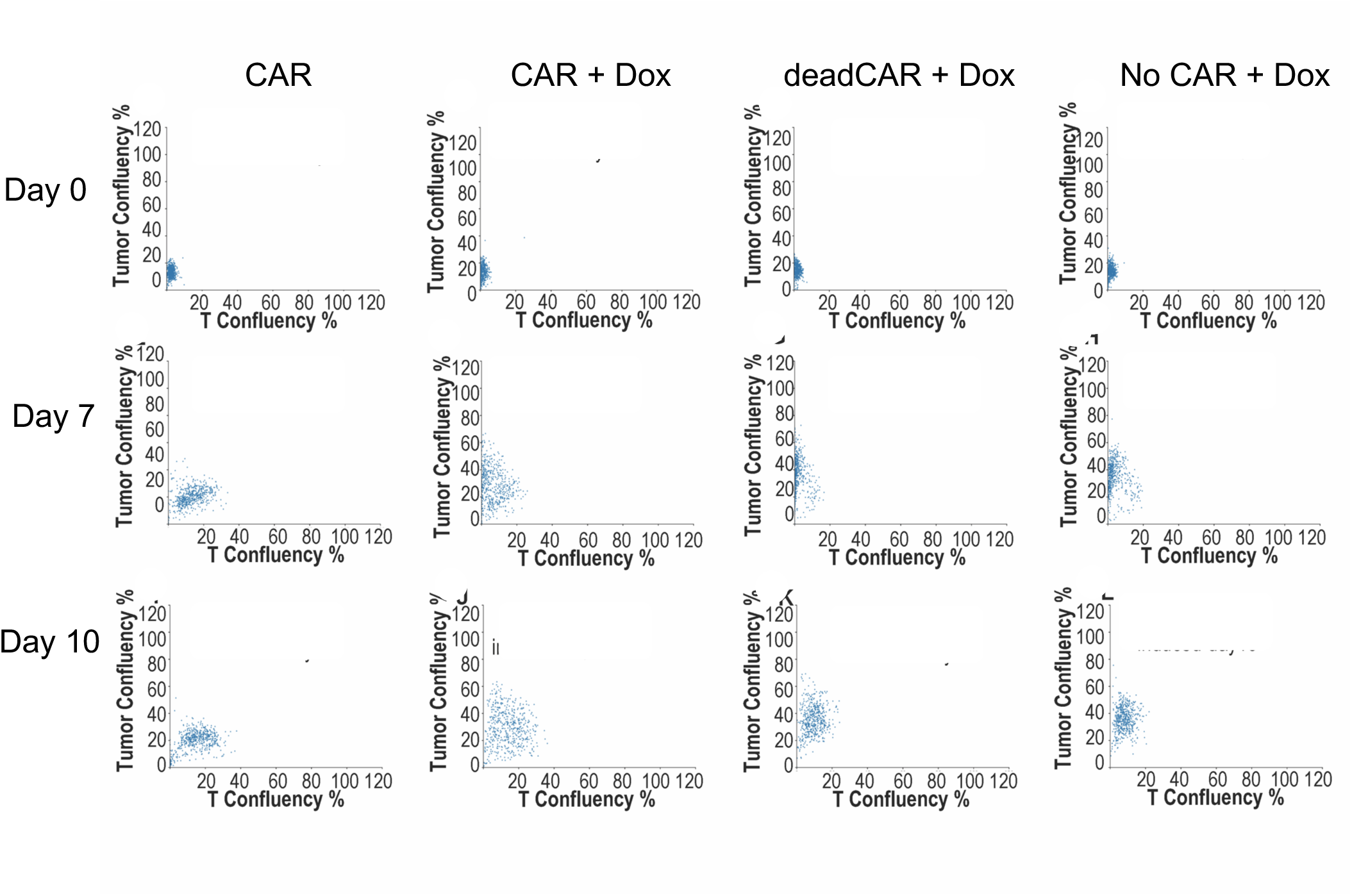
Evolution of T cell mediated cytotoxicity in individual microwells. On the key dates of initial coculture (Day0), second feed (Day7) and third feed (Day 10), The T cell confluency(x) and Tumor cell confluency(y) of each individual microwells were plotted as scatter plot of the ∼500 microwell of each for inducible-CD19-CAR uninduced, inducible-CD19-CAR induced, inducible-CD19 dead-CAR induced, inducible-CD19 no CAR induced.

